# A Fully Implantable Wireless Bidirectional Neuromodulation System for Mice

**DOI:** 10.1101/2021.06.02.446797

**Authors:** Jason Wright, Jason Wong, Jose Mathew, Ibrahim Mughrabi, Naveen Jayaprakash, Theodoros P. Zanos, Yousef Al Abed, Stavros Zanos, Timir Datta-Chaudhuri

## Abstract

Novel research in the field of bioelectronic medicine requires systems that pair high-performance neurostimulation and bio-signal acquisition hardware with advanced software signal processing and control algorithms. Although mice are the most commonly used animal in medical research, the size, weight, and power requirements of such systems either preclude their use or impose significant constraints on experimental design. Here, we describe a fully-implantable neuromodulation system suitable for use in mice, measuring 2.2 cm^3^ and weighing 2.8 g. A bidirectional wireless interface allows simultaneous readout of multiple physiological signals and complete control over stimulation parameters, and a wirelessly rechargeable battery provides a lifetime of up to 5 days on a single charge. The device was successfully implanted (N=12) and a functional neural interface (capable of inducing acute bradycardia) is demonstrated with functional lifetimes exceeding to three weeks. The design utilizes only commercially-available components and 3D-printed packaging, with the goal of accelerating discovery and translation of future bioelectronic therapeutics.

## Introduction

The emerging field of bioelectronic medicine aims to utilize electrical interfaces to the nervous system to modulate immune and metabolic state. Unlike for drug discovery, there are few standardized tools and approaches for developing and validating novel therapies for electroceutical approaches. Though there are a myriad of disease models in mice, new bioelectronic medicine studies require the development of specialized technologies and preclinical models because most neuromodulation devices are too large for use in mice. In this work we discuss the development of a fully implantable generalizable stimulation and sensing platform for implantation in mice, enabling the development of new therapies using standard disease models and validation techniques. Though neuromodulation technology as a whole has advanced considerably in recent years, developing systems for interfacing with the nervous system of mice has a number of critical challenges associated with the size of the animal and the corresponding constraints on mass, volume, material choices, and the mechanical challenges of interfacing to delicate nerve bundles. To overcome these challenges requires the use of miniaturized electronics development of reliable chronic electrodes, non-standard biocompatible packaging, and wireless telemetry and power.

Current methods for chronic neuromodulation in mice require the use of tethered systems, preventing normal behavior, introducing stress due to handling, and in the case of anesthesia, suppressing the body’s response to interventions. A robust generalizable platform for chronic wireless stimulation and monitoring in freely moving mice does not exist. In addition to electrical stimulation and recording, such systems should be capable of monitoring motion and temperature, and accommodate future modular sensor integration by incorporating additional signal conditioning circuits. Such a platform will facilitate previously impossible research by allowing stimulation in response to physiological and neural signals over biologically relevant time periods permitting the study of disease and immune response progression.

### Opportunities for bidirectional systems

Typical medical interventions have a separation of therapy and assessment approaches, meaning that a different method is used to deliver an intervention than to assess its efficacy. A simple example is administration of a pharmacological agent, and a blood test to determine the effect of the drug. Bioelectronic medicine approaches utilize electrical interfaces to the nervous system to deliver stimulation-based therapy in the same manner as existing active implantable devices such as pacemakers and deep brain stimulation systems. Much of the information about the immune and physiological state of the body is also conveyed by the nervous system, and as such, there is an opportunity for bioelectronic medicine devices to also assess the efficacy of an intervention at the same interface at which that intervention is applied. This opportunity points to the utility of bidirectional implantable devices for the development of new bioelectronic therapies. In addition to monitoring electrical activity in the body, long-term bidirectional implants can also monitor other changes over time such as biomolecule concentrations, blood pressure, and physical activity.

Closed-loop systems can further build upon the capabilities of bidirectional systems by incorporating therapy modulation based on tracked effectiveness over time. This allows for tuning of therapies for individual subjects, and also over the course of disease progression and recovery. The need for closed-loop approaches in clinical applications has long been recognized [1-3]. Recent indications under active research include spinal cord injury [4, 5], psychiatric disorders and neurodegenerative diseases [6, 7], bladder dysfunction [8], diabetes management [9], hypertension [10], and pain relief [11]. In some applications, the closed-loop algorithm is relatively simple, e.g. enabling or disabling a pre-set stimulation paradigm based on a physiological measurement being above or below a pre-determined threshold. Other applications require more complex and computationally demanding algorithms. Closed-loop systems may also benefit from partitioned algorithms that are divided into classifiers (state estimation) implemented on an embedded device and control policy (state update) implemented offline [12]. This approach simplifies the demand for on-board computation while retaining advanced functionality.

### Device development for mice

Medical research is primarily performed in mice due to the availability of a diverse set of disease models and the maturity of genetic and pharmacological approaches for developing new models. While closed-loop neuromodulation devices have been developed and used in pre-clinical research for many years, no existing solutions provide a bidirectional neural interface in a single implantable form factor suitable for use as a generalizable closed-loop system for mice. The size of implants for mice is limited by the mass and volume that they can support. While there is no universal standardized guidance for limits on implantable devices for mice, typical Institutional Animal Care and Use Committee (IACUC) guidelines advise that mouse models with tumor limit the size of single tumors to not exceed 20mm in their largest dimension, and to stay at or near 10% of body weight [13]. Commercially available mouse telemetry devices (primarily from Data Sciences International) range in mass up to 2.2 gm, and have volumes up to 1.7 cm^3^, these devices are known to be well tolerated for implant durations of a few months. Additionally, implants must be constructed from biocompatible materials to minimize the risk of infection, tissue necrosis, and/or an inflammatory foreign body response.

In [14] and [15], a fully implantable device meets all requirements, but its packaged volume and mass make it unsuitable for use in mice. In [16], a wireless neural recording system is demonstrated in freely moving mice, but it does not contain electrical stimulation capabilities and is not implantable. In both [8] and [17], a flexible implant demonstrates wirelessly controllable optogenetic stimulation, but without biosignal recording functionality. To date, there are a number of commercial devices for implantation in mice that meet the size and mass constraints, but have specialized functionality such as glucose or blood pressure sensing, or limited biopotential recording capability such as electrocardiography (ECG) or electromyography (EMG). None of these existing devices provide a general-purpose platform for bidirectional neuromodulation in mice.

In this work, we present the first wireless, fully implantable bidirectional general-purpose neuromodulation system suitable for chronic use in mice, utilizing only commercial off-the-shelf (COTS) components, 3D-printed packaging, and commercial electrodes for ease of manufacturability. At 2.2 cm^3^ and 2.8 g, this system can be used to achieve an untethered neural interface for several weeks, while providing sensor interfaces and computational capability needed for future closed-loop experiments. Chronic *in vivo* testing demonstrates functional neural interface lifetimes beyond 3 weeks.

## Results

### System architecture

Figure 1 shows the overall system architecture of the bidirectional neuromodulation system described in this work. The system consists of an implanted device, a holding cage equipped with a wireless charging transmitter, and an external PC. The implanted device provides on-board computation, a neural recording and stimulation controller, a front-end for ECG sensing, an inertial measurement unit, auxiliary amplifiers for future sensor integration, a bidirectional wireless interface for telemetry and configuration, and a wirelessly rechargeable battery. The device is fabricated on a 6-layer FR4 PCB measuring approximately 18mm by 15mm, along with a flexible polyimide printed circuit board for resonant power transfer matching the same outline, and was packaged using a 3D printed shell enclosure prior to implantation. Details of the different system components are provided in the materials and methods section.

**Figure 1.**
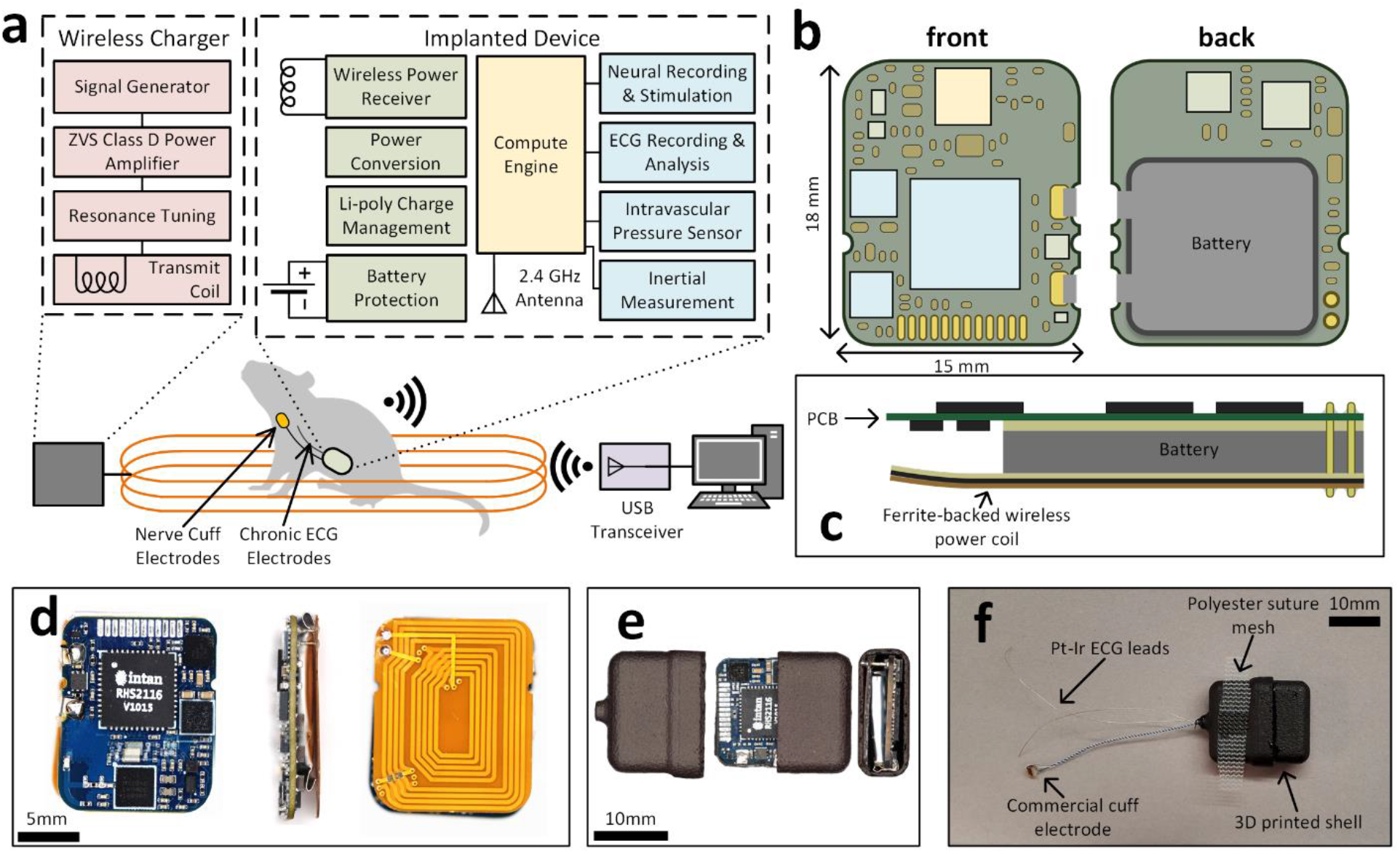
(a) system diagram showing functionality and communication between the implant and host PC, (b) PCB layout showing relative component sizes and dimensions, (c) diagram of assembled device stackup, (d) photographs of assembled electronics, (e) the electronics contained within the two-part 3D-printed shell enclosure, (f) final device with an attached nerve electrode, ECG leads, and suture mesh ready for implantation.

### Electrical stimulation and bio-potential sensing

In this work, a 16-channel neural stimulation and recording engine (RHS2116, Intan Technologies, Los Angeles, CA, USA) is utilized for its high degree of functionality in a single package, and a long history of successful use in neuromodulation applications. The RHS2116 has independent source-sink current DACs and programmable bandwidth amplifiers for each channel. For this application, the stimulation capabilities of the RHS2116 were desired for several reasons. First, the specifications are comfortably inclusive of typical stimulation parameters in mice: amplitudes from 10 µA-2.5 mA [18], frequencies from DC-1 kHz, and pulse widths from 10 µs-500 µs [19]. Second, the ability to accurately program an arbitrary stimulation waveform is essential for many applications. For example, stimulation of the cervical vagus nerve in mice may either increase or decrease production of certain cytokines such as tumor necrosis factor (TNF) and interleukin-10 (IL-10), depending on the combination of amplitude, frequency, and pulse width used [20]. Non-rectangular waveforms can have a differential impact on fiber recruitment and can be used to reduce overall charge injection and energy consumption [21, 22]. Finally, the RHS2116 can provide both biphasic stimulation and passive charge balancing of the electrode, which is essential for both safety considerations [23] and for maintaining a functional electrode interface [24].

The RHS2116 also meets typical requirements for neural recording in mice. A sample rate of at least 20 ksps is needed to accurately capture action potentials (APs), which have a duration of approximately 2 ms and occur at a rate of up to 500 Hz [25]. Compound action potentials (CAPs), which consist of a weighted aggregate of individual Aps, are more likely to be measure from a peripheral nerve interface and have similar characteristics. These “spikes” of neural activity typically have an amplitude in the low µVs (for example, baseline recordings of the vagus nerve in mice can show amplitudes less than 15 µVpp [26]).

For ECG recording, a single-chip analog frontend with on-chip R-to-R interval computation (MAX30003, Maxim Integrated, San Jose, CA, USA) is utilized. The MAX30003 is available in a very small wafer-level package (8 mm^2^) and requires very few external components. Although its onboard algorithms are intended for use in human ECG systems, the parameters can be adjusted to support the differing qualities of the ECG signal in mice. The MAX30003 offers a lower-power recording interface for ECG signals for which the higher performance of the RHS2116 is not required.

An on-board microcontroller with integrated 2.4 GHz transceiver (nRF52840, Nordic Semiconductor, Trondheim, Norway) is used to control sensor interfaces and provide a bidirectional wireless interface to the host PC. While some applications may not require a host PC (for example, on/off stimulation control based on measured HR being above/below a pre-programmed threshold), others require computationally expensive signal processing techniques, for example to mitigate the impact of stimulus artifacts on neural recordings, where a combination of front-end and back-end signal processing techniques provides the best result [27]. In such applications, the round-trip latency between the occurrence of a neural event and subsequent stimulator output is of particular interest. An ideal system can respond to single-neuron spikes with minimum latency. In applications such as motor prostheses (involving restoration of sensory proprioception), epilepsy management and seizure control [28], and plasticity [29], low latencies are required to achieve a desired effect, ideally in the µs to low ms range. In this work, the total round-trip latency (measured as the time required for the implant to send a packet of data, the host PC to receive that packet and send a response, and for the implant to receive the response packet) is typically 450-500 µs.

For future modular sensor integration, several additional circuits are included: a Wheatstone bridge circuit is used to excite the piezoresistive pressure sensor and its differential voltage is amplified and converted to a single-ended signal using an integrated instrumentation amplifier (AD8235, Analog Devices, Wilmington, MA, USA), again chosen primarily for small package size (3.2 mm^2^), and a 3-axis MEMS accelerometer is utilized (LIS3DH, STMicroelectronics, Geneva, Switzerland) from which pose and activity level can be estimated using an offline algorithm, although these sensors were not utilized in this work. A more detailed depiction of this work’s sensing and stimulation circuits is shown in Supplementary Figure 1.

#### Stimulation

Performance of the stimulation circuit was assessed using a resistive load (500 Ω, the standard test condition for nerve and muscle stimulators per IEC 60601-2-10). The results for a range of test conditions similar to those expected for chronic stimulation of a mouse vagus nerve are shown in **Error! Reference source not found**.(a-b). Representative stimulation output is shown for 4 different amplitudes of symmetric charge balanced biphasic waveforms. The integral nonlinearity (INL) of output current, a measure of the deviation from ideal, was calculated for currents between -200 µA and 200 µA.

#### ECG recording

To evaluate performance of the ECG recording circuit, a reference dataset of mouse ECG recordings from PhysioZoo [30-32] was used as the input to a signal generator (National Instruments USB-6341) configured with differential output and connected to the ECG inputs on the device. The signal generator DAC has a precision of ∼305 µV resulting in some distortion to the signal. The recordings were streamed wirelessly with ECG gain set to 160V/V and sample rate set to 512 sps. Peaks of the R wave were counted over a 1500ms moving window (threshold of 1.5 mV and 50ms spacing) to calculate the beats per minute (BPM). An example result using the first 10 seconds of Mouse01 in the PhysioZoo dataset is shown in **Error! Reference source not found**.(c).

### In vivo experiments

#### Acute validation

All animal experiments were conducted with the approval of the Institutional Animal Care and Use Committee at the Feinstein Institutes for Medical Research. A female C57BL/6 mouse was anesthetized with isoflurane and a cuff electrode placed on the cervical branch of the vagus nerve. Identical stimulation parameters were programmed into our device and a commercial stimulator (STG-4000, MultiChannel Systems GmbH, Reutlingen, Germany) and delivered to the nerve to assess the effect on heart rate (Bio Amp/PowerLab, ADInstruments, Sydney, Australia). Brief durations of stimulation correspond to bradycardia; the relationship between stimulation amplitude and magnitude of bradycardia is compared between the two systems in Figure 3(a) and (b). In a separate procedure under identical conditions, one channel of the device was used to record from the cuff electrode. ECG and respiratory EMG signals were clearly visible (corresponding to a heart rate of ∼450 BPM and a respiratory rate of ∼55 BPM), with spontaneous neural activity with amplitude of about 100 µV occasionally manifesting in the signal.

**Figure 2:**
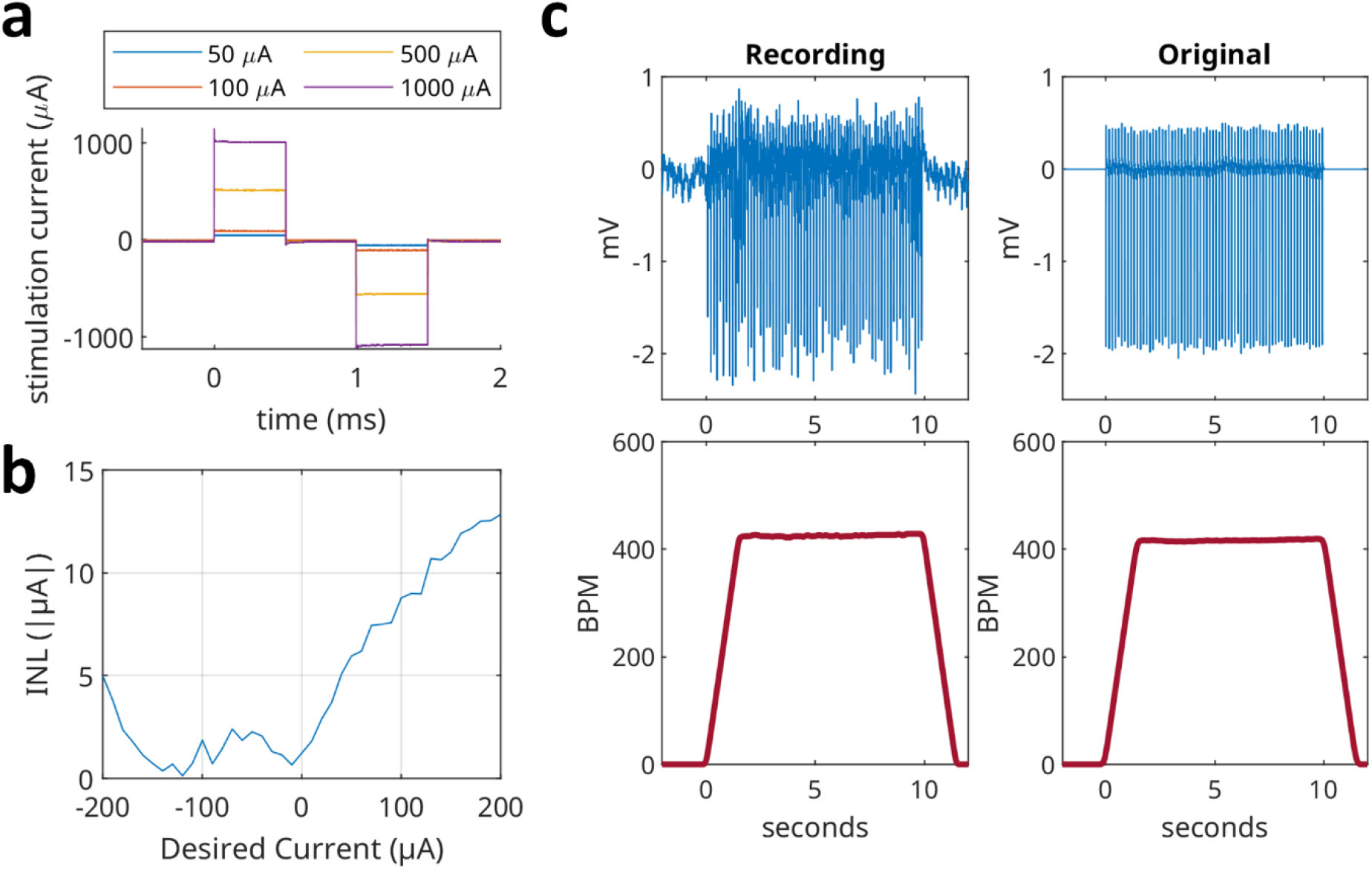
(a) representative stimulator outputs as measured across a 500 Ω load with pulse widths and inter-pulse interval set to 500 µs, (b) integral non-linearity for the stimulator output in the ±200 µA range. (c) comparison between wireless stream of ECG data (left) and original recording as input to signal generator (right), using offline BPM calculation.

**Figure 3.**
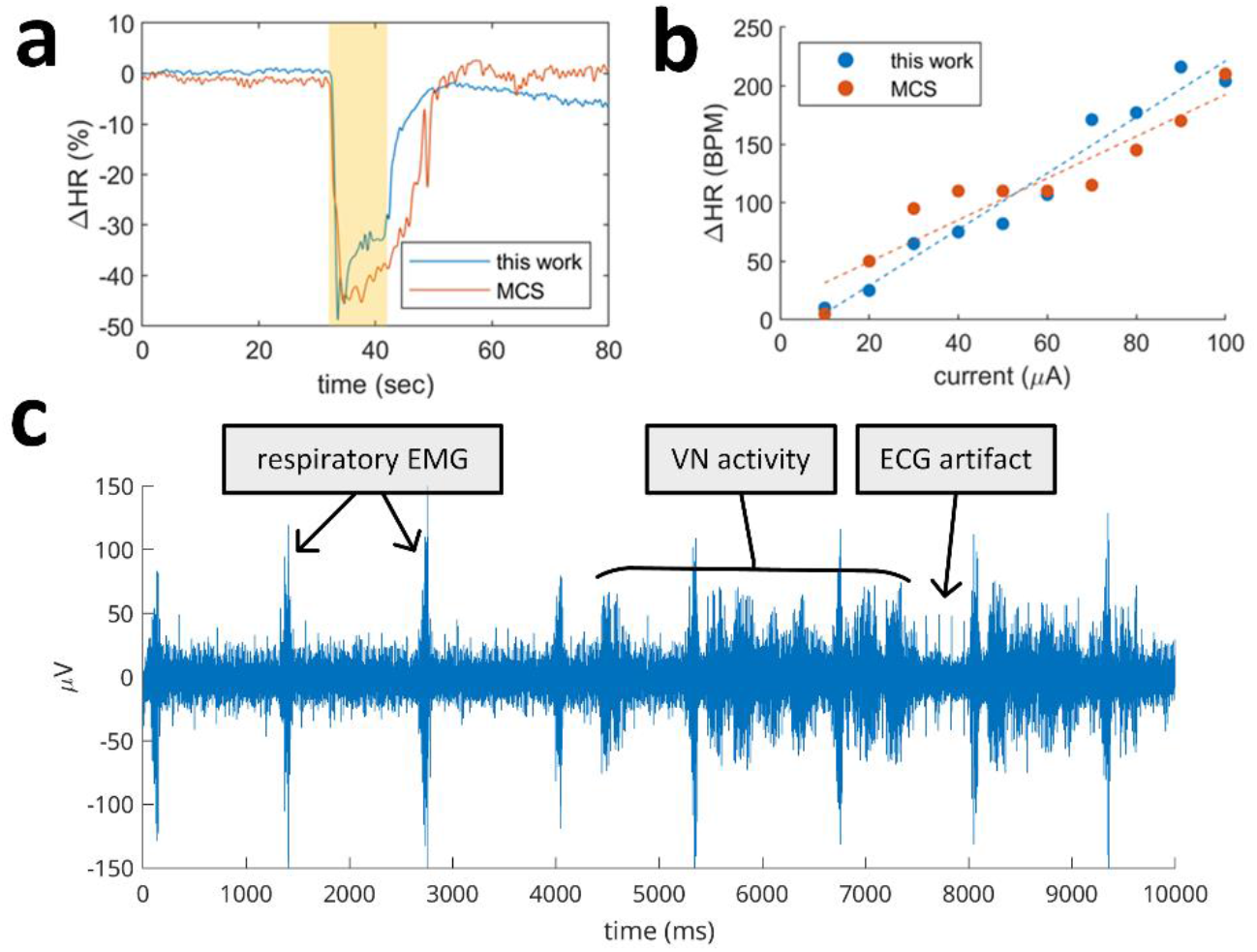
(a,b) Functional equivalence testing indicating that the stimulation performance of the system is equivalent to a commercial benchtop stimulator, evaluated by single 10 second stimulation trains (a), highlighted in yellow for the period in which stimulation is active, and by a comparison of the amplitude dependence of induced bradycardia (b). (c) Representative recording showing artifacts from ECG and respiratory EMG interspersed with spontaneous VN activity.

#### Chronic implantation

A total of 11 devices were implanted chronically using either an intra-abdominal or dorsal subcutaneous approach into C57BL/6 mice. Of the 11 devices, 1 was re-implanted after explanation, for a total of 12 implantation procedures. All of the mice survived the implantation procedure. Stimulation was performed immediately following implantation (day 0) to determine if the device and nerve cuff interface were functional. The devices were tested at one day intervals (starting at day 1 post-implantation) for up to a week after implantation, and approximately two times per week after that to determine successful operation in a chronic setting.

To evaluate implant functionality, brief (2-10 second) periods of 30 Hz stimulation were used with pulse widths ranging from 100-1000 µs and amplitudes ranging from 5-2500 µA. Implants were tested with subjects under anesthesia and in awake, freely moving subjects. ECG recordings were captured in both cases; for anesthetized animals ECG recordings were also performed using the ADInstruments system described previously. Figure 4(a) shows an example of stimulation-induced bradycardia under anesthesia shortly after implantation, validated by using the external system for comparison. Figure 4(b) shows an example of the same effect persisting in the same animal over 1 week post-surgery.

**Figure 4.**
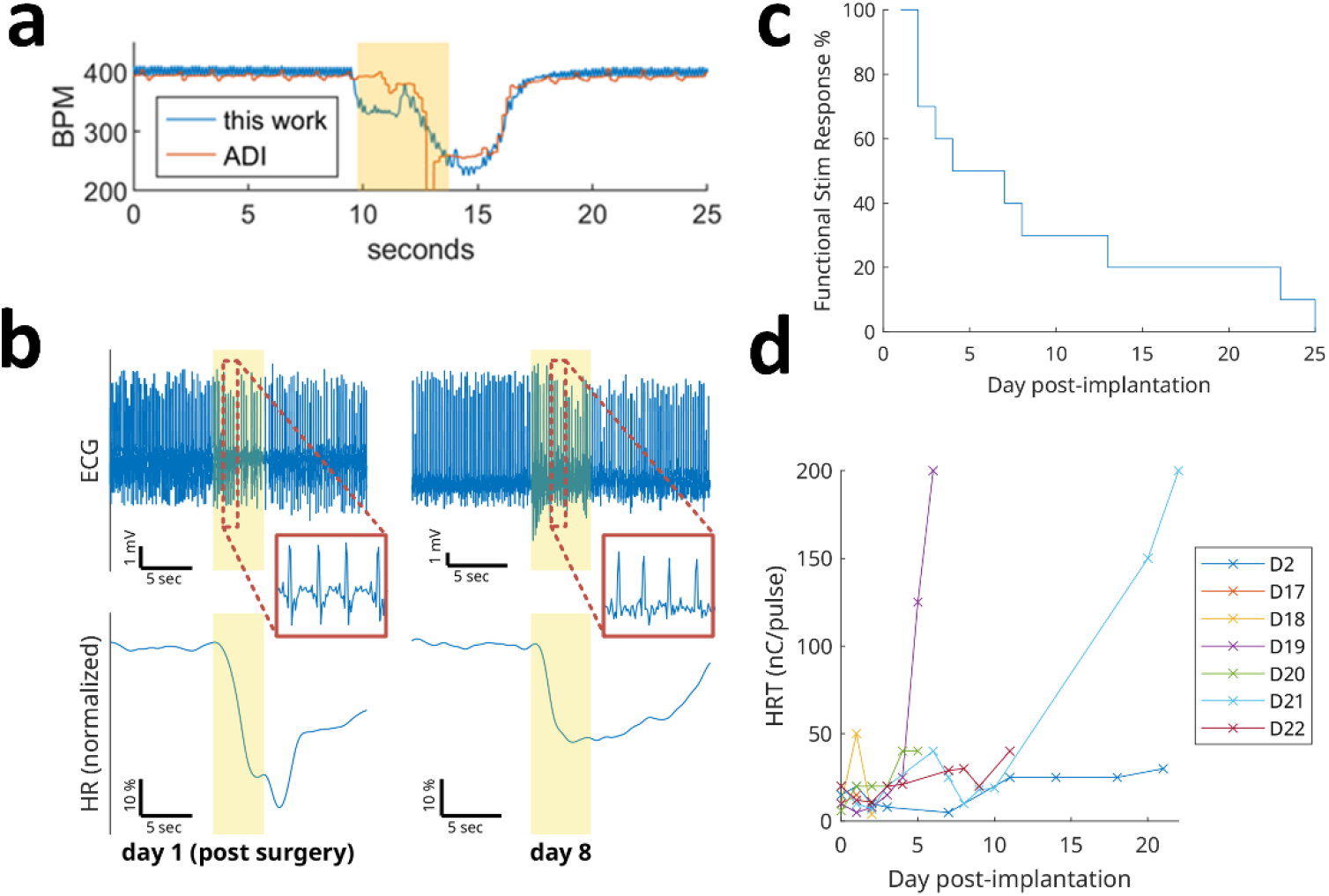
(a) Day 0 ECG recording (top) and comparison between implant (on-board) recordings and external ADI system (bottom). Stimulation was active during the highlighted region. In this example, BPM was computed offline. The stimulation parameters used were 10 µA amplitude, 100 µs pulse width, 30 Hz frequency. (b) Example of stimulation-induced bradycardia reproduced over a week post-implantation in the same animal. In this example, the implant ECG recording is used to compute BPM offline. Stimulation parameters used were 100µA/100µs/30Hz on day 1 and 250µA/100µs/30Hz on day 8. This testing was performed under anesthesia. (c) Percentage of implants showing a functional stimulation response over time, defined as a 5-15% drop in HR following stimulation. (d) HRT for each implant over time.

Of the 12 implantations, 2 devices failed to elicit a response on day 0, and 3 devices failed to elicit a response on day 1. Lifetime statistics were calculated using the remaining 7 implantations that continued to produce a heart rate response to stimulation for at least day 1 after implantation. All of the 7 devices functioned electronically, as determined by successful wireless communication and charging until explantation, which in some cases was beyond the point at which they no longer produced a response to stimulation. The device survival curves are shown below in Figure 4(c), where “survival” is defined as capable of inducing a ∼5-15% heart rate drop in response to stimulation. Many devices were able to induce muscle contractions in response to stimulation, even when there was no reduction in heart rate, persisting beyond one month. All of the mice survived until the (terminal) explantation procedure, and no devices failed from a packaging or electronics perspective.

Due to electrode interface degradation and/or fibrosis stemming from chronic cuffing, the intensity of stimulation required to achieve a ∼5-15% drop in heart rate tends to gradually increase over the lifetime of an implant [33]. This heart rate threshold (HRT) was continually measured for each implant showing a response beyond day 1, and the results are shown in Figure 4(d) as a function of total charge per pulse.

### Power and Telemetry

#### Circuit power consumption

Power consumption of the device was measured in various operational modes using an external supply (Keithley SourceMeter 2400, Cleveland, OH, USA) set to 3.7V output. The idle power consumption was measured as 85 µW, and representative results for an active configuration are shown in Figure 5(a). Idle power consumption includes the microcontroller waking from sleep and transmitting a packet once per second. These results indicate a battery life of about 5 days (idle) or 35 minutes (active, most features enabled).

**Figure 5.**
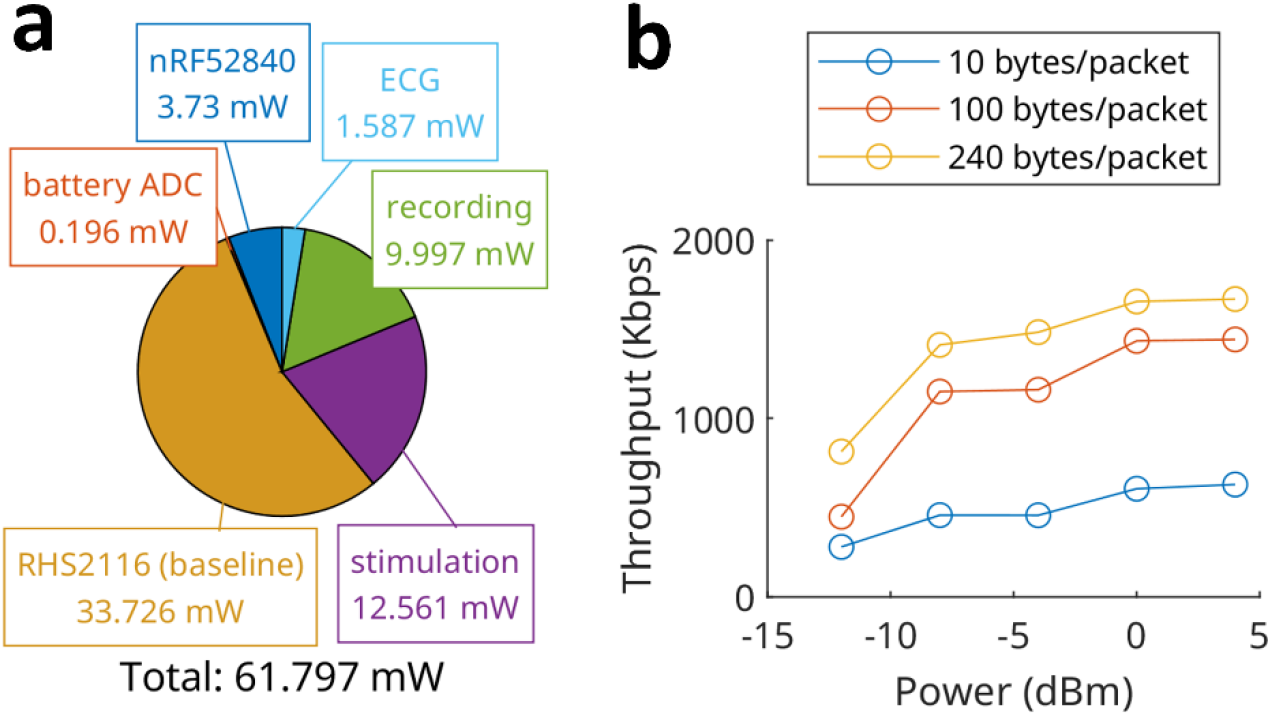
(a) power consumption of the entire system in active (most features enabled) mode. The radio output power was set to +4 dBm. Stimulation parameters used were biphasic ±100 μA, 100 μs pulse width, 30 Hz frequency, no load, 1 channel. Neural recording was configured as 1 channel at 20 ksps. (b) effect of varying output power levels and packet lengths at a distance of 30cm (unpackaged PCB) averaged over a 10 second interval.

#### Wireless control and telemetry

Bidirectional wireless communication is needed to receive data generated by the implant (telemetry) and for external control of the implant’s state. The telemetry throughput should generally be at least 320 kbps (the rate needed to support a continuous 20 ksps stream of neural recordings with 16 bits per sample), although this requirement could be relaxed if compression, event filtering, or other techniques are used [34]. Because mice are housed in relatively small environments (typical cage dimensions are roughly 15-30 cm), range is not critical. The receiver can be placed anywhere outside the cage or even inside the cage to maintain a minimum distance of less than 30cm at all times. Throughput was assessed by varying the output power setting and packet length per transmission while keeping a fixed distance between device and transceiver. As shown in Figure 5(b), maximum throughput can be achieved by maximizing output power and packet length, at the cost of increased power consumption and increased sample latency.

### Wireless Charging

#### Powering implanted devices

Numerous techniques exist for powering implantable electronics. They may be broadly grouped into battery-powered and battery-free systems, such as ultrasonic, optical, piezoelectric, or chemical energy harvesting devices. For this work, a secondary (rechargeable) battery-powered approach was used, for two main reasons: first, a closed-loop device with high-bandwidth transmission exceeds the power limitations of many battery-free power delivery techniques; second, these techniques may interfere with other parameters of the experiment, such as by constraining animal motion, requiring the animal to be outfitted with additional hardware, or by generating electromagnetic noise that interferes with sensitive recordings.

Although primary (non-rechargeable) batteries offer the highest energy density, the power requirements of the system described so far cannot be met with currently available primary cells given the volume and mass constraints for implantation in mice, and so a secondary battery is required. The most common technique for recharging implanted electronics is inductive power transfer, requiring the target device to be close to the source transmitter. While this approach works well for current clinical embodiments, it limits the free movement of animal subjects (as power may not be available during rearing or other behaviors) and can bias the outcome of experiments when the animal is handled or restrained.

The efficiency of wireless power systems varies widely depending on the design of the receiver coil and the transmitter-receiver separation and alignment, most neural implants are limited to a power budget of <100 mW at a maximum distance of 25mm when utilizing near-field inductive coupling [35]. In this work, only about 40 mW is usable for battery charging (limited by the maximum safe charge current). We developed a custom wireless power system cable of delivering at least 40 mW throughout a standard mouse cage, with minimal variation to coil orientation and an efficiency of 0.58% at the edge of the cage and 0.38% at the center (defined as the efficiency of the entire system, including losses from the power amplifier, AC/DC converter and battery charger).

#### Magnetic resonant coupling

Most commercial wireless power applications (e.g. mobile phone charging) use inductive coupling and require the receiver to be placed very close to the transmitter at a particular orientation. For a chronic implant in a freely moving mouse, a wider freedom of position and orientation is required. As a result, wireless power systems for small animal implants often utilize magnetic resonant coupling [36, 37], a mid-range technique that utilizes resonance to maximize efficiency between loosely-coupled coils. In this work, the wireless power system follows a conventional 4-coil approach, with a source coil and an intermediate resonant transfer coil on the transmit side, and an intermediate resonant receiver coil and a load coil on the implant side [38]. The transmit coils were wrapped around an enclosure matching the dimensions of a standard mouse cage, with the source coil connected to the supply and the resonant coil isolated electrically from other components. The system is depicted in Figure 6(a,b).

**Figure 6.**
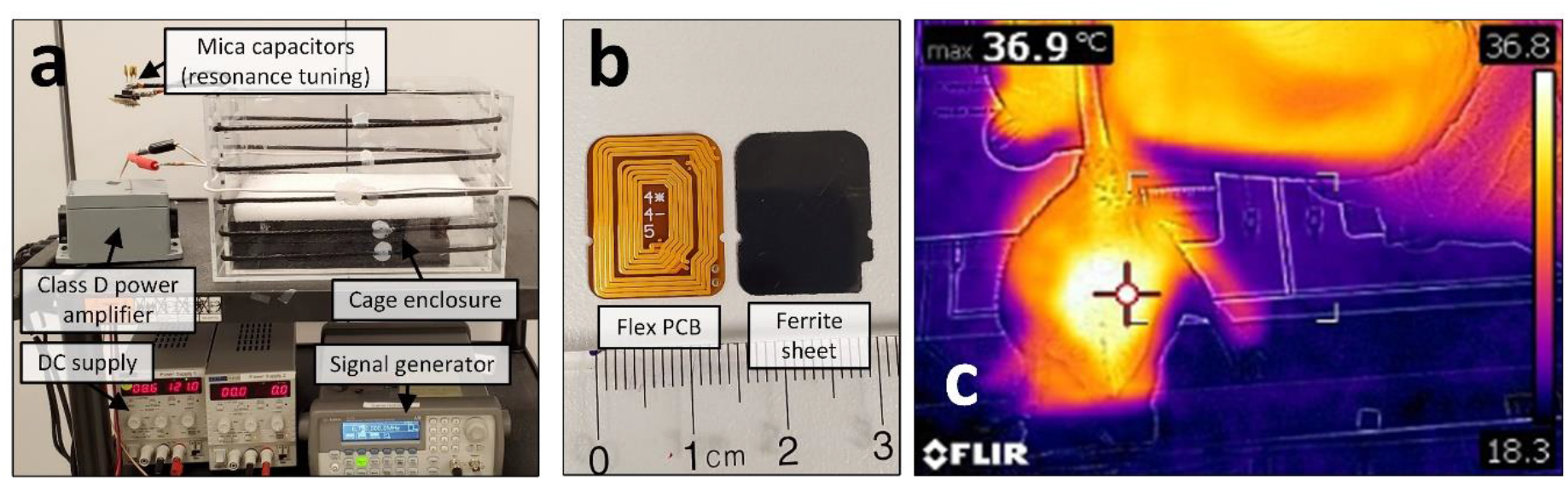
(a) Wireless power transmitter setup with cage enclosure, power amplifier and signal source, (b) flex PCB coil and 300µm ferrite sheet used for assembly, (c) thermal image of the mouse with the device implanted after 2 hours of charging.

#### Safety considerations

Any wireless power transmission system necessitates important safety considerations. Wireless power transfer is a form of radiated energy and as such there are limitations on the levels of exposure which are considered safe. In the United States, continuous exposure to electromagnetic fields is limited to a specific absorption rate (SAR) of 1.6 W/kg (human head model) to avoid negative health effects from tissue heating; this acts as an upper limit on transmitted power. For implanted medical devices, 2 °C is considered the upper bound of safe chronic temperature increases, corresponding to a power density of approximately 40 mW/cm^2^ to 80 mW/cm^2^ [39, 40].

Due to circuit power dissipation and use of wireless charging, both the implant and surrounding tissue are at risk of temperature increase during operation of the implant. Several benchtop experiments were conducted prior to in vivo experiments to ensure animal safety, using both representative tissue samples and packaged implants (detailed in Methods). Figure 6(c) shows a thermal image of the abdomen with the implanted device after charging for 2 hours. During all animal experiments, the mouse grimace scale was used to assess pain; at no point did any animal show signs of severe pain or discomfort.

#### Implant power management

A full-bridge rectifier with fast Schottky barrier diodes and an integrated voltage regulator (XCM414, Torex Semiconductor, Tokyo, Japan) converts the MHz signal induced on the receiver coil to a DC voltage. This part was chosen due to its small package size (7.54 mm^2^), high input voltage tolerance (26V) and protection afforded to downstream circuits. The 5V output variant of this part was chosen to provide a suitable input to typical lithium polymer battery chargers which generally require at least a 4.2V supply. An integrated lithium polymer charger (BQ25101, Texas Instruments, Dallas, TX, USA) provides the appropriate constant current/constant voltage (CC-CV) charging profile, with its charge limit and termination programmable via external resistors. A protection circuit (BQ29700, Texas Instruments, Dallas, TX, USA) monitors the battery voltage and current and disconnects the cell from the rest of the circuit in the event of various fault conditions. In particular, under-voltage lockout (UVLO) and overvoltage protection (OVP) ensure the battery is not damaged by either condition via breakdown of electrode materials, dendrite formation, or overheating, minimizing the risk of damage to the implant and/or the animal. An overview of the power management circuit is shown in Supplementary Figure 2.

### Packaging

The neuromodulation system must be packaged in order to protect the electronics from the harsh implant environment while creating a biocompatible surface suitable for contact with the surrounding tissue. The packaging shields the device from moisture and ions which can cause device failure by creating electrical shorts, causing corrosion, or result in other electrochemical interactions leading to electronics failure [41, 42]. Packaging materials must be biocompatible in order to have an implant that is well tolerated by the animal. Use of materials that are not biocompatible or not properly processed or handled can lead to a wide range of adverse effects that can jeopardize the animal’s health and/or impact the experiment. Other factors such as mass, volume and impact on wireless performance (where applicable) are also critical to the design of implantable electronics packaging.

In this work, a custom 3D-printed shell package consisting of two interlocking halves (as shown in Figure 1) was used. Shells were fabricated with 500 and 750 µm wall thickness, coated with a 20 µm layer of parylene C, and filled with 300 mg of silica gel desiccant. After placing the electronics into the shells, the two halves and the electrode feedthroughs were sealed with epoxy and cured.

To evaluate packaging efficacy, custom humidity-sensing boards utilizing 3 different commercially available humidity sensors (SHTW2, Sensirion AG, Stäfa, Switzerland; HDC2010, Texas Instruments, Dallas, TX, USA; HTS221, STMicroelectronics, Geneva, Switzerland) were built and packaged using the same approach. After sealing with epoxy, the packaged humidity sensors were fully immersed in 1X PBS and stored in an oven at 60 °C. Humidity data was regularly collected from all sensors. As shown in Figure 7, humidity inside the packaging reached the 50% mark at around three weeks, with the 750 µm wall thickness offering only a marginal improvement.

**Figure 7.**
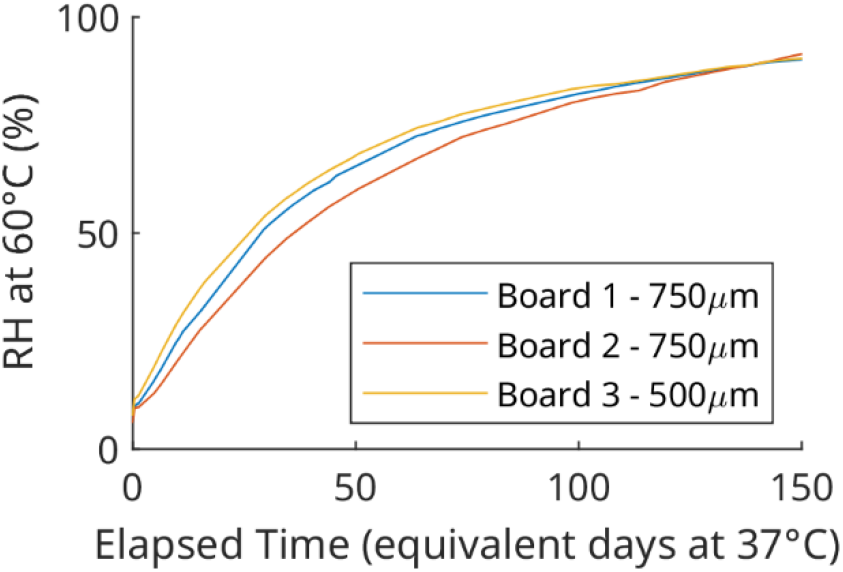
Aggregate humidity data for accelerated-life-tested devices in shell packaging. Plots show an average of readings from the three different sensors for each board. Arrhenius equation is used to compute the equivalent lifetime at mouse body temperature of 37 °C.

## Discussion

We have demonstrated the first bidirectional neuromodulation system with a sufficiently small volume and mass for chronic implantation in mice. Integration of the widely-used Intan RHS2116 neuromodulation frontend with modern commercially-available integrated circuits enables a system small enough and with a sufficiently low power budget for continuous fully-implanted operation in short- to medium-term chronic experiments. Designing a device that can be easily manufactured (relying only on COTS components and 3D-printed packaging) was a priority, as many efforts in related areas do not see broader adoption among researchers due to the cost and complexity of producing devices. This work was performed using two different commercially available cuff electrodes, showing that stimulation and sensing electrodes can be independently selected and subsequently integrated with this system.

Table 1 summarizes the system specifications in comparison to other neuromodulation systems that demonstrate either capabilities similarly suitable for closed-loop operation (including on-board computational power and fully wireless interfaces), a small fully-implantable form factor, or a combination of the two. All systems demonstrate both technical performance and *in vivo* validation.

**Table 1.** Comparison to selected neuromodulation systems.

| System | NeuroChip-2 [43] | PennBMBI [44] | U. Toronto [45] | DSI HD-X series | Neuropace RNS | Activa PC+S [46] | WAND [47] | Bionode [14] | WiNERS-8 Implant [15] | This Work |
| --- | --- | --- | --- | --- | --- | --- | --- | --- | --- | --- |
| Dimensions (mm) | 63 × 63 × 30 | 56 × 36 × 13 (rec.), 43 × 27 × 8 (stim.) | 22 × 30 × 15 | 20 × 10.75 × 8.9 | 28 × 60 × 7.7 | 65 × 49 | 36 × 33 × 15 | 35 × ø15 | 30 × 15 × 5 | 19.9 × 18.1 × 6.6 |
| Volume (cm<sup>3</sup>) | 119.07 | 35.48 | 9.9 | 1.4-1.7 | 12.93 | 39 | 17.82 | 6.19 | 2.25 | 2.22 |
| Weight (g) | 145 |  | 12 | 2.2 | 16 | 67 | 17.9 | 9 | 2.8 | 2.8 (incl. 0.3g desiccant) |
| Power (mW) | 284-420 | 290 | 45 | -- | -- | -- | 172 | ~10-50 | 18.9 | 62 |
| Wireless | IR | ESB | ZigBee | 455 kHz | Inductive | 175 kHz | BLE | 2.4 GHz | 433 MHz | ESB |
| Data rate (Mbps) | 0.024 | 2 | 0.25 | -- | -- | 0.011 | 1.96 | -- | 9 | 2 |
| Real time | No | 4 ch | 1 ch | 3-4 | 1 ch | 2 ch (raw)<br>4 ch (compr.) | 96 ch + 3 accel ch | 2 ch | 32 ch | 8 ch |
| # Stim. channels | 3 | 2 | 64 | 0 | 8 | 8 | 128 | 1 | 4 | 4 |
| # Recording channels | 3 | 4 | 256 | 3-4 | 4 | 4 | 128 | 2 | 32 | 8 |
| Sampling rate (kS/s) | 24 | 21 | 15 | -- | 0.25 | 0.422 | 1 | 25 | 25 | 20 |
| ADC resolution (bits) | 8 | 12 | 8 | -- | 10 | 10 | 15 | 8 or 10 | 10 | 16 |
| Max. Stim. Current (mA) | 5 | 1 | 0.24 | -- | 11.5 | 25.5 | 5 | 1.125 | 1.86 | 2.5 |
| Stim. Compliance (V) | ±15 | ±12 | 2.6 | -- | 12 | ±10 | 12 | ±10.5 | ±2 | +6/-12 or +12/-6 |
| Compute power | 2x M8C | AVR | -- | -- | -- | -- | Cortex-M3 | Cortex-M0 | None | Cortex-M4 |
| Animal model | Primate | Rat | Rat | Mouse | Human | Ovine | Primate | Rat | Rat | Mouse |

### Opportunities for translation and future work

This work demonstrates a system that intends to advance the technological path towards the fully-instrumented “wireless mouse” as a frontier for bioelectronic medicine research. Many previous discoveries have not been translatable to real-world therapeutics at least in part because large-scale studies in freely-moving animals are impractical or impossible. As development, manufacturing, implantation, and experimentation with this system advances, research involving large numbers of subjects becomes possible. A fully implanted system allows testing to be done with virtually zero animal handling and no limitations on subject mobility due to tethering. A charger- and transceiver-equipped cage could automatically conduct routine experiments with minimal human intervention. Future work will aim to further reduce the device size and weight, demonstrate autonomous closed-loop operation, and utilize remaining computational power to reduce the amount of data that needs to be transferred by the implant.

## Methods

### Software

#### Firmware

The implanted device runs custom firmware written in C, built using the nRF5 Software Development Kit

(SDK). The firmware is responsible for controlling the RHS2116 and other sensing circuits, condensing acquired data into ESB packets, and receiving and applying state updates generated by the host PC. Further details are available in Supplementary Information (Supplementary Figure 3).

#### Custom USB transceiver

To collect data from the implanted device, off-the-shelf development kits were used (PCA10056/PCA10059, Nordic Semiconductor, Trondheim, Norway). Custom firmware was written to send and receive packets to/from the implanted device, converting the data from the wireless format to USB packets. The transceiver uses the Communications Device Class (CDC) Abstract Control Model (ACM) profile to enumerate as a standard serial port on the host PC. Escape characters are used to delimit packets.

#### Graphical user interface

Custom software was developed using the Qt C++ framework to provide a graphical user interface (GUI) to the system operator. The GUI allows reading/writing of the implanted device’s state as well as a live view of incoming data streams during an experiment. The software can also write this data to a file for later analysis. A real time R-to-R interval algorithm is also implemented to allow heart rate computation from a raw data stream, as an alternative to the on-chip algorithm provided by the MAX30003. A representative screenshot is shown in Supplementary Figure 4.

### Wireless telemetry

#### Physical layer

The ISM 2.4 GHz band is chosen for wireless telemetry to minimize antenna size, minimize attenuation due to tissue absorption, and maximize bandwidth. Although the ISM 2.4 GHz band is not ideal given the typical dielectric properties of biological tissue [48], it allows the use of a highly miniaturized antenna suitable for our implant’s size constraints while supporting the high bandwidth needed to support real-time neural recording telemetry. A ceramic chip antenna (SRCW004, Antenova Ltd, Hertfordshire, UK) provides an extremely compact solution. While our implant size and component density does not allow for the standard conductor keep-out requirements for a chip antenna (typically several mm on all sides), we are able to tolerate reduced performance due to our short range requirement.

#### Data layer

The system uses the Enhanced ShockBurst (ESB) protocol developed by Nordic Semiconductor. This protocol provides useful features such as CRC integrity checks, packet acknowledgement, and automatic retransmission of packets. In comparison to Bluetooth Low Energy (BLE), ESB can be used for very low latency applications by avoiding the minimum connection interval (CI) of 7.5ms mandated by the BLE specification. The implanted device uses the PTX (transmitter) mode and the external transceiver uses the PRX (receiver) mode to minimize power consumption on the implanted device side, as the radio can be turned off when not in use only in PTX mode. Further details on the ESB implementation are available in Supplementary Information (Supplementary Figure 5).

### Power

#### Battery

Lithium ion batteries are widely available and offer high volumetric efficiency and high peak discharge currents, and as a result are commonly used in implantable neurostimulators [49]. For this work, GEB201212 (PowerStream Technology, Orem, UT, USA; General Electronics Technology Shenzhen Co., Ltd, Shenzhen, China) was used, primarily due to its small size (12mm × 12mm), high capacity (10 mAh), high discharge rate (1C), and low mass (0.5g).

#### Operating frequency selection

Most inductive systems operate at kHz frequency (for example, commercially available Qi chargers, which operate at 80-300 kHz), but resonant systems are operated at a higher frequency in MHz range. For this application, the operating frequency must lie between an upper limit imposed by tissue absorption and a lower limit depending on signal bandwidth and the need to avoid radio interference on the medium waveband [50]. For this reason, a frequency of 6.78MHz was selected.

#### Wireless power transfer and coil design

The wireless power system in this work is based on [38] using a 4-coil strongly-coupled resonant system, consisting of a source coil, a transmit resonant coil, a receive resonant coil and a load coil. The source coil and the load coil are connected directly to the source and the load respectively and the remaining two coils are tuned to the operating frequency of 6.78 MHz using a suitable capacitor connected in parallel. To maximize the quality factor of a coil, it is optimized for a combination of high self-inductance and low series resistance [51, 52].

Transmit coils were constructed using 44AWG/2400 strands Litz wire (Cooner Wire, Chatsworth, CA, USA) to minimize the impact of skin effect and wrapped in a helical manner around a rectangular acrylic sheet, sized to fit a standard mouse cage as seen in Supplementary Figure 7(c). Receiver coils were fabricated on a 4-layer 1oz flexible PCB of size 18mm × 15mm and thickness of 300µm. The flexible PCB construction allows for minimum thickness, but with the drawback that deformation of the flex coil during assembly causes a shift in inductance and thereby the resonant frequency, although careful assembly can mitigate this problem. Supplementary Figure 6(a) shows the shift in resonant frequency of an example device after each assembly step; tuning the transmit coil to match the resonant shift on the receiver is used to compensate for this shift [53]. Parasitic capacitance of the resonant coil was controlled by adjusting the space between turns; multiple designs were fabricated and tested to find the optimum coil design. Due to the placement of the coil parallel to the aluminum battery pouch, an eddy current is induced which alters the inductance and detunes the resonance of the LC circuit. Placing a magnetically active ferrite sheet helps to shield unwanted eddy currents, thereby increasing the transfer efficiency [54]. 300 μm ferrite sheets (MARUWA Co., Owariasahi, Japan) were laser-cut to size and affixed to each coil.

#### Power amplifier

The transmit coil is driven by a Class D eGaN FET based zero-voltage-switching (ZVS) amplifier [55]. The GaN based FET offers low input and output capacitance and low inductances, allowing it to operate at high efficiency at the intended frequency. Here we used EPC2007C GaN transistors (EPC, El Segundo, CA, USA), which have low capacitance and low on resistance (R_DS(on)_) at desired operating frequency. Using MOSFETs requires higher gate charge than eGaN FETs, which incurs higher gate driver power losses and reduced efficiency [56]. Apart from using a simple class E amplifier which is popular for wireless power transfer, a class D amplifier was used because the class E amplifier is sensitive to load variation [57]. This provides ease of charging the implanted device by regulating the input power to the transmit coil without performance loss in the amplifier. The power amplifier board is kept inside an enclosed box with cooling fan and heat sink. Figure 6(a) shows the wireless power transfer cage enclosure connected to the EPC power amplifier fed by a MHz square wave using a signal generator and DC power supply.

#### Thermal considerations

To evaluate heating caused by the wireless power system, representative tissue samples were placed inside the charging cage at the edge where maximum magnetic field exposure is seen and monitored for temperature increases and compared to a control sample. Supplementary Figure 6(b) shows the change in temperature of the tissue sample after exposing the magnetic field continuously for more than two hours. No increase beyond 1.5 °C was detected after 90 minutes of continuous exposure on the tissue sample (transmit power set to 8W). Additionally, an experiment to measure the temperature of the implant itself was conducted. Two samples each of unpackaged (initial) devices, packaged devices, and packaged devices immersed in PBS were prepared. The transmit power level was adjusted to deliver a constant 5mA or 10mA charge to the battery for each device. The results are shown in Supplementary Figure 6(c). Of note, both tests show a sharp temperature rise corresponding to the point at which the battery charging circuit switches from constant-current to constant-voltage output (after about 120 minutes at 5 mA and about 60 minutes at 10 mA). During in vivo experiments, the battery voltage was monitored to avoid reaching the CV transition point.

### Assembly, packaging, and cleaning

PCBs were manufactured using lead-free RoHS-compliant processes. Prior to packaging, devices were cleaned ultrasonically using multiple cycles of deionized water and isopropyl alcohol to remove any surface contaminants that may have formed during manufacturing, shipping, or handling, as such contaminants can negatively impact implant lifetimes through mechanisms such as dendrite formation or corrosion.

The shell package consists of two 3D-printed shell halves conformally coated with parylene C. The shells enclose the PCB assembly and are sealed with biocompatible epoxy (M31-CL, Loctite, Düsseldorf, Germany). During the assembly process, available space in the package cavity is filled with silica gel desiccant to reduce the rate of humidity increase. The electronics packaging was designed to conform closely to the shape of the PCB assembly to minimize total implant volume. Shells were printed in HP 3D High Reusability PA 12 material (HP Inc., Palo Alto, CA, USA), selected for its biocompatibility as well as its fluid tightness [58]. The parylene layer acts as an additional barrier, creating a hydrophobic surface with reduced pore size. Surface hydrophobization provides a small benefit to packaging effectiveness by limiting water adsorption and surface transport. Reducing surface pore size increases the length of the diffusion pathway into the device and as a result slows transmission rate of permeants [59].

Immediately prior to implantation, devices were sterilized via submersion in Cidex OPA solution (Advanced Sterilization Products Inc., Irvine, CA, USA).

### Electrodes and surgical technique

Individual cuff electrodes from Micro-Leads (Somerville, MA, USA) and CorTec GmbH (Freiburg, Germany) were connected to one channel of each device prior to packaging. Co-coiled leads were used to provide flexible lead length. ECG leads were prepared from PtIr wire and trimmed to size for each implant. Implants were placed in either the abdomen or the back and secured with suture mesh. Supplementary Figure 7(a,b) shows the surgical procedure and subcutaneous lead tunneling for an abdominal implant.

## Acknowledgements

Funding for this work was provided by Northwell Health.

