## Supplemental Information for "A Fully Implantable Wireless Bidirectional Neuromodulation System for Mice"

### Supplementary Information

#### Hardware

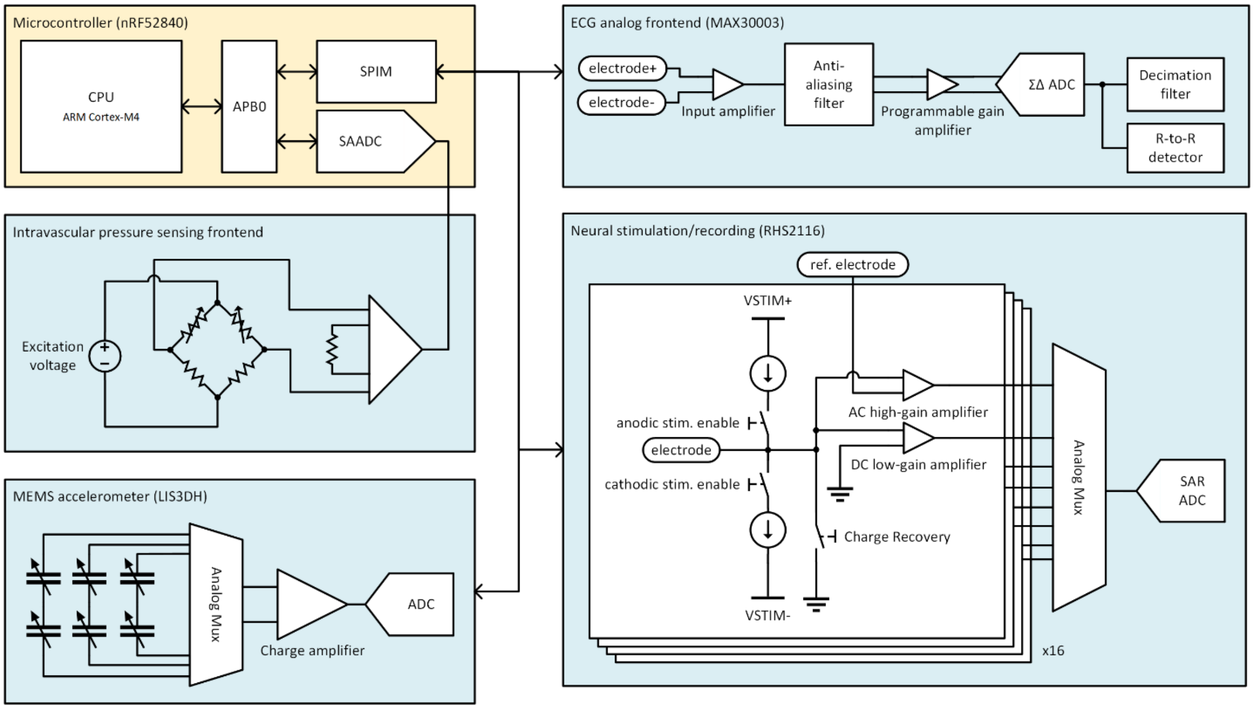


*Supplementary Figure 1. Diagram of the stimulation and sensing components of the design and their interface with the microcontroller.*

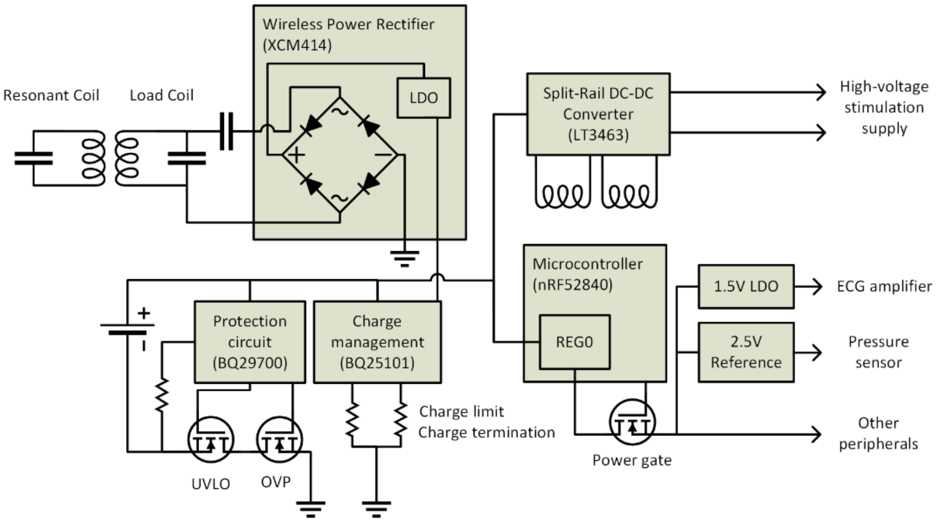


*Supplementary Figure 2. Architecture of the implanted device power management. Wireless power received by the resonant coil is converted to DC and distributed to the battery charging circuits, the high voltage power supply for stimulation, and to the on-board microcontroller which provides power to the other peripherals in the system.*

**Software**

*Firmware architecture*

Firmware is structured around the producer-consumer pattern. Various modules ("producers"), each representing a sensor, regularly generate data and push it to a buffer of samples. Each sample is tagged with a timestamp and the sensing channel from which it originated. Another module (the "consumer") monitors the buffer of samples and converts them to packets to be streamed wirelessly. A double buffering is used to separate memory for incoming and outgoing samples to ensure new samples can always be handled. Each module can be enabled/disabled or configured according to the system state. Supplementary Figure 3 shows a high-level view of the firmware architecture. Most operations are triggered off of a system timer (10 kHz), while a separate timer with higher priority is used for higher sample rate recordings (20 kHz). Recorded samples are processed via interrupt service routine. The main thread is reserved for managing the data buffer and queueing wireless packets as needed.

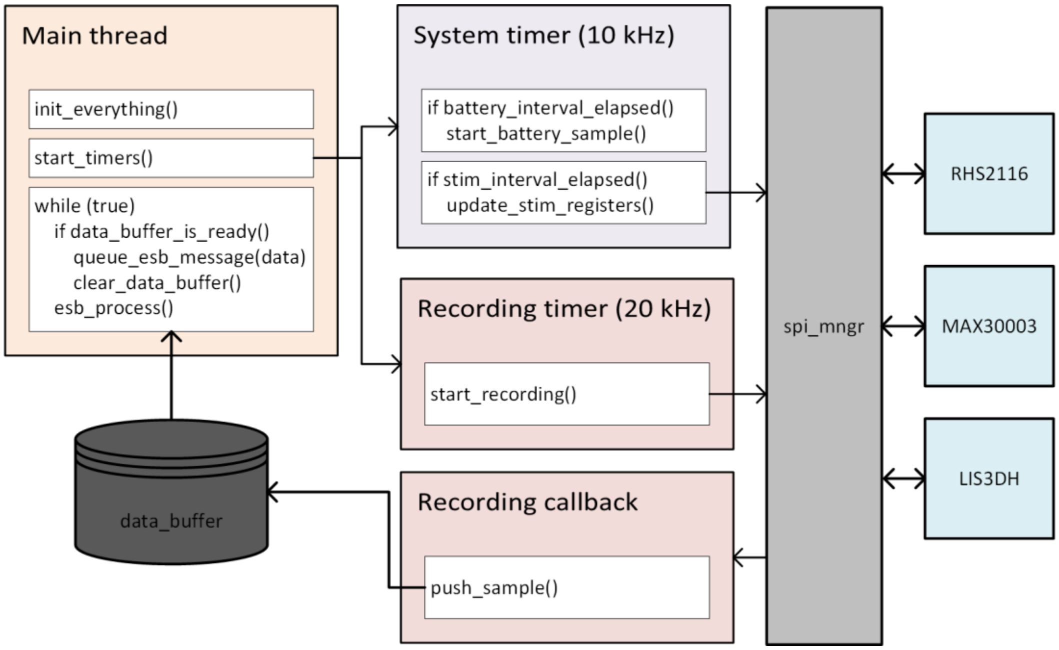


*Supplementary Figure 3. High level (pseudocode) view of firmware operation.*

*PC GUI*

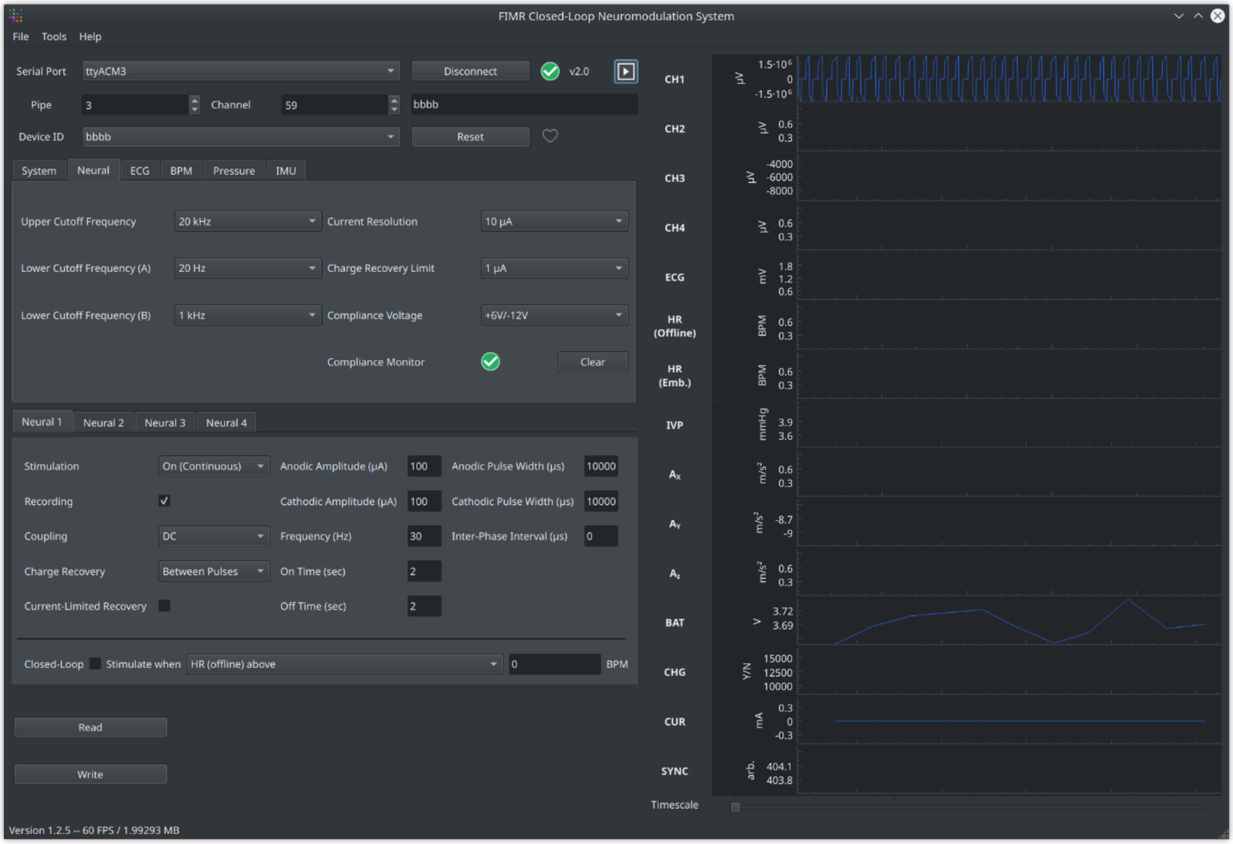


*Supplementary Figure 4. Screenshot of the custom GUI for controlling the implanted device.*

*ESB implementation*

Each ESB packet is formatted as shown in Supplementary Figure 5.

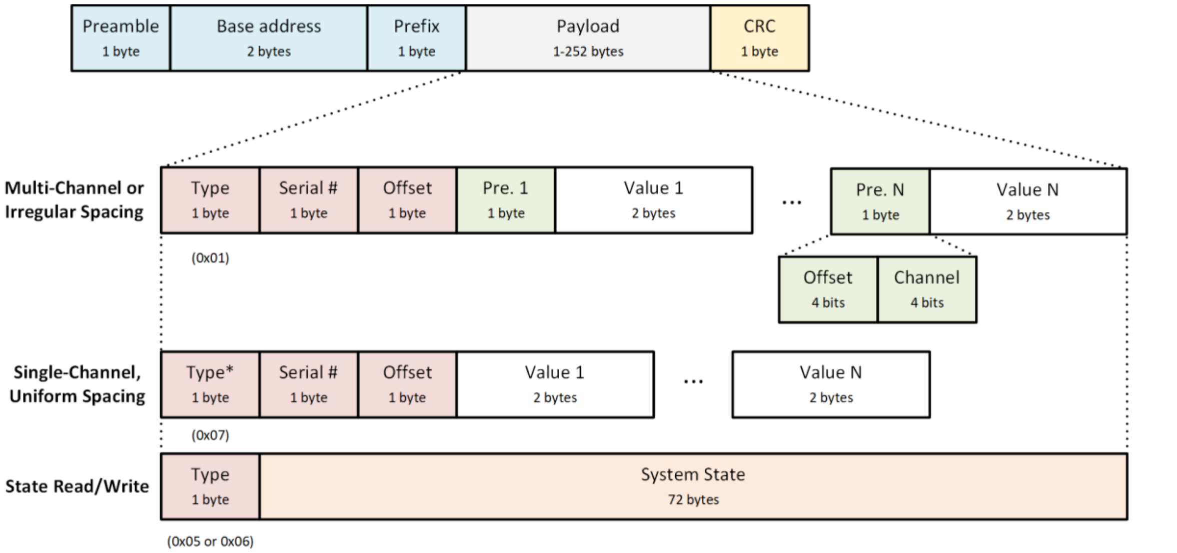


*Supplementary Figure 5. ESB packet format, showing example payload formats for three different packet types. *For the single-channel case, the Type byte has its upper four bits set to the channel ID used for all samples in that packet.*

Most ESB traffic consists of either (a) new samples being sent from the implanted device to the host, or (b) a new system state being written to the implanted device, e.g. to update stimulation parameters. For (a), two different packet layouts are used depending on the contents of the sample buffer. If samples from multiple channels are present, or if the channel sampling is not uniform, the first layout is used. Otherwise, the second layout is used, offering slightly more efficiency by avoiding the preamble byte (labeled "Pre" in Supplementary Figure 5) which encodes the time offset for each sample and the channel ID. However, other packet types are used (the remaining 4-bit values of the "Type" byte) to communicate errors, partial updates, and for testing and debugging.

**Wireless power**

It was found that 4 turns per layer in a 4 layer board had the highest Q factor given the coil dimensions and operating frequency. The trace width was chosen such that the resistance is minimized and to allow space for the other (load) coil in the same planar structure. Supplementary Table 1 shows the characteristics of the transmit and receive coil measured using Keysight E5061B VNA at 7 MHz.

| | **Transmit coil** | | **Receiver Coil** | |
|---|---|---|---|---|
| | Source coil | Resonant coil | Resonant coil | Load coil |
| Turns | 1 | 5 | 12 | 15 |
| Inductance (μH) | 1.34 | 11.9 | 4.22 | 2.33 |
| Series resistance (Ω) | 0.396 | 12 | 9.6 | 5.2 |
| Self-resonant frequency (MHz) | 30 | 14 | 18 | 30 |

*Supplementary Table 1. Parameters for custom designed transmitter and receiver coils.*

To resonate at 6.78 MHz, the transmit resonant coil has a 63pF capacitor connected in parallel and a 75pF capacitor for the receiver resonant coil.

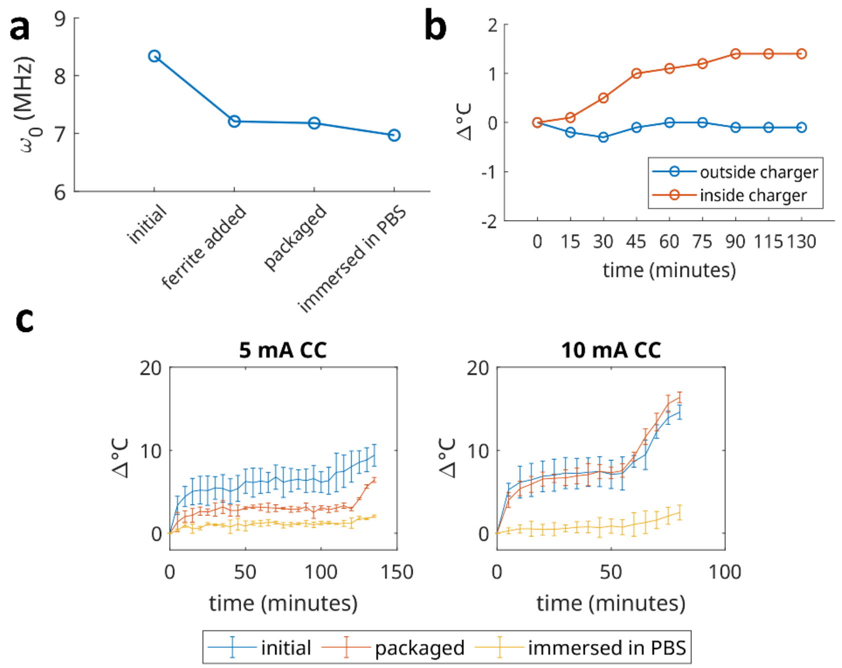


*Supplementary Figure 6. (a) resonant frequency shift during each stage of assembly, (b) tissue heating of sample under maximum magnetic field exposure, (c) temperature rise of the implant due to wireless charging at two different power levels.*

#### In vivo experiments

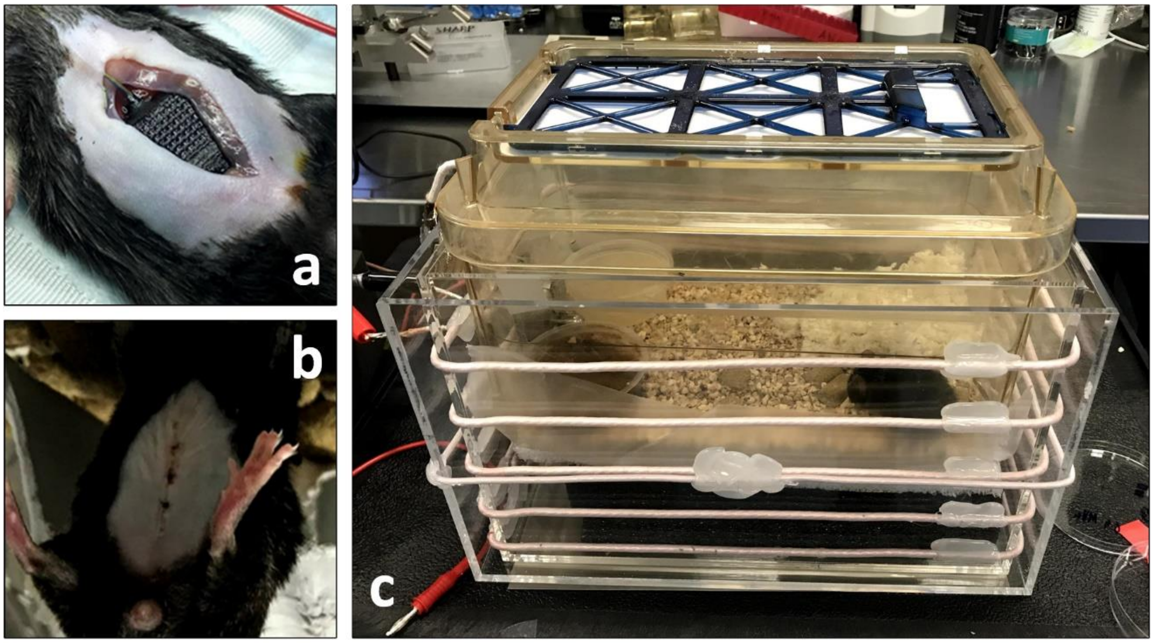


*Supplementary Figure 7. (a) implant being placed intra-abdominally with subcutaneous tunneling of electrode leads, (b) surgical site of the same animal 7 days post-implantation, (c) freely moving animal in wireless charging cage.*
